# Male bumblebees (*Bombus terrestris*) are more explorative and behaviourally flexible than workers

**DOI:** 10.1101/2025.03.28.645856

**Authors:** Pizza Ka Yee Chow, Sophie Donelly, Kevin D. Hochard, Theo Robert

**Author notes:** Corresponding author:* Pizza Chow.

## Abstract

Sex role differences can influence ecological and evolutionarily important traits, such as exploration and behavioural flexibility. In bumblebees, for instance, females (workers) are the main foragers for the colony whereas males (drones) have minimal responsibility. When males leave the nest, however, they become solitary foragers while searching for mates for reproduction. This suggests that increased exploration and behavioural flexibility are crucial for their survival and reproduction success. Here, we compared the exploratory behaviour and behavioural flexibility of male bumblebees (*B. terrestris*) with that of female workers in a laboratory. We measured bees’ active time in a novel environment in an exploration task and their colour-reward associative learning ability and behavioural flexibility in a simultaneous two-choice discrimination-reversal colour learning task. As predicted, males were more active (thus explored more) in the novel environment than females. Performance of a colour-reward association learning was comparable between males and females, but males demonstrated enhanced behavioural flexibility when the reward contingency changed. Males used the ‘win-stay, lose-shift’ strategy, making fewer errors (being more flexible) than females. Males’ explorative behaviour may be related to their pre-mating patrolling behaviour in the wild. Their enhanced flexibility suggests their readiness to find profitable flowers when other flowers decrease in quality. These results highlight the importance of exploration and behavioural flexibility for males, which can increase their chance of attracting mates for reproduction and maximising their energy during foraging. Results provide insights into how different traits are related to sex roles in various fitness-related contexts.

## Introduction

Behavioural and cognitive traits, such as exploration and behavioural flexibility, are linked to fitness [1,2]. Exploration allows individuals to discover resources and gain information about their environment [3]. Behavioural flexibility allows individuals to adjust their behaviours according to environmental demands and opportunities [4,5]. The expression of these traits may vary within a species depending on the role of sexes in different fitness-related contexts (e.g., [6–9]). Investigating such variation is crucial for understanding the ultimate significance of these traits.

A classical model for sex role difference is the bumblebee, *Bombus spp*.. During the colony development, females (workers) of bumblebee species, such as (*B. terrestris*), are the main foragers and carers for the broods. Males (drones) emerge in the mid to end of the colony cycle, do not forage for the colony as they lack pollen baskets [10], and have minimal responsibility before dispersal [11–15]. When drones leave their nest, they rarely return home [14,16]. They start solitary foraging and looking for mates for reproduction. Compared with females, males’ cognitive and behavioural traits remain largely unexplored but the significant differences in sex roles may lead to variations in ecologically and evolutionarily important traits, such as exploration and behavioural flexibility, between females and males.

Males generally fly longer and travel farther than workers [17–21], which may benefit males’ exploration of their environment and increase their chances of finding an unrelated queen for reproduction and discovering profitable flower patches [17,22]. Traits like associative learning of profitable flower colour and location have been shown in males [16,23–27], and behavioural flexibility is expected to be a crucial trait for solitary males to maximise their energy gain and improve their survival chances, such as adapting their behaviour after a flower quality degrades. Here, we designed an ecologically relevant experiment to measure the active time (a proxy for exploration) of male and female bumblebees in a new environment with an exploration task. Their associative learning ability and behavioural flexibility were also tested in a traditional two-colour discrimination-reversal learning task [29]. We predicted that males would show more exploratory behaviour and higher behavioural flexibility than females.

## Methods and Results

### Study species and housing

Bumblebees from 5 colonies were individually marked with a numbered tag using shellac-made natural glue [30] (details in Note S1). Bees were kept in a black box, mirroring their underground nest. The nest was attached to a transparent circular tunnel (diameter 4 cm), which had three shutters and an open end (to capture a bee using a plunger for the experiment). The lab (room temperature 21-23^°^c) was lit with natural daylight and artificial lights. The bees had *ad libitum* access to syrup and pollen in the exploration task but were supplied with a decreased amount of syrup and pollen during the learning task to increase their motivation. This study obtained ethics approval (Note S2).

### Sex and Exploration in a New Environment

Sixty-nine bees (33 females, 36 males) left their nest for the first time and participated in the exploration task during which they could freely move in ‘the exploration box’. This box was divided into 10 equal-sized compartments with the ‘floor’ of it covered with a random white-and-red checkerboard pattern (figure 1A, Note S3). A bee could pass between compartments through a hole in the middle of the divider. Their time spent being active in each compartment of this new environment was measured as exploration [31], which was recorded by a video camera hung above the box. For each bee, we summed up the active time across compartments (i.e., add up the time spent in every compartment) to obtain the total active time. Each bee was fed with 50 w/w sucrose *ad libitum* [32–34] when s/he exited the box (left the last compartment). We recorded the amount of sucrose consumption as a proxy of sucrose responsiveness/hunger level.

**Figure 1.**
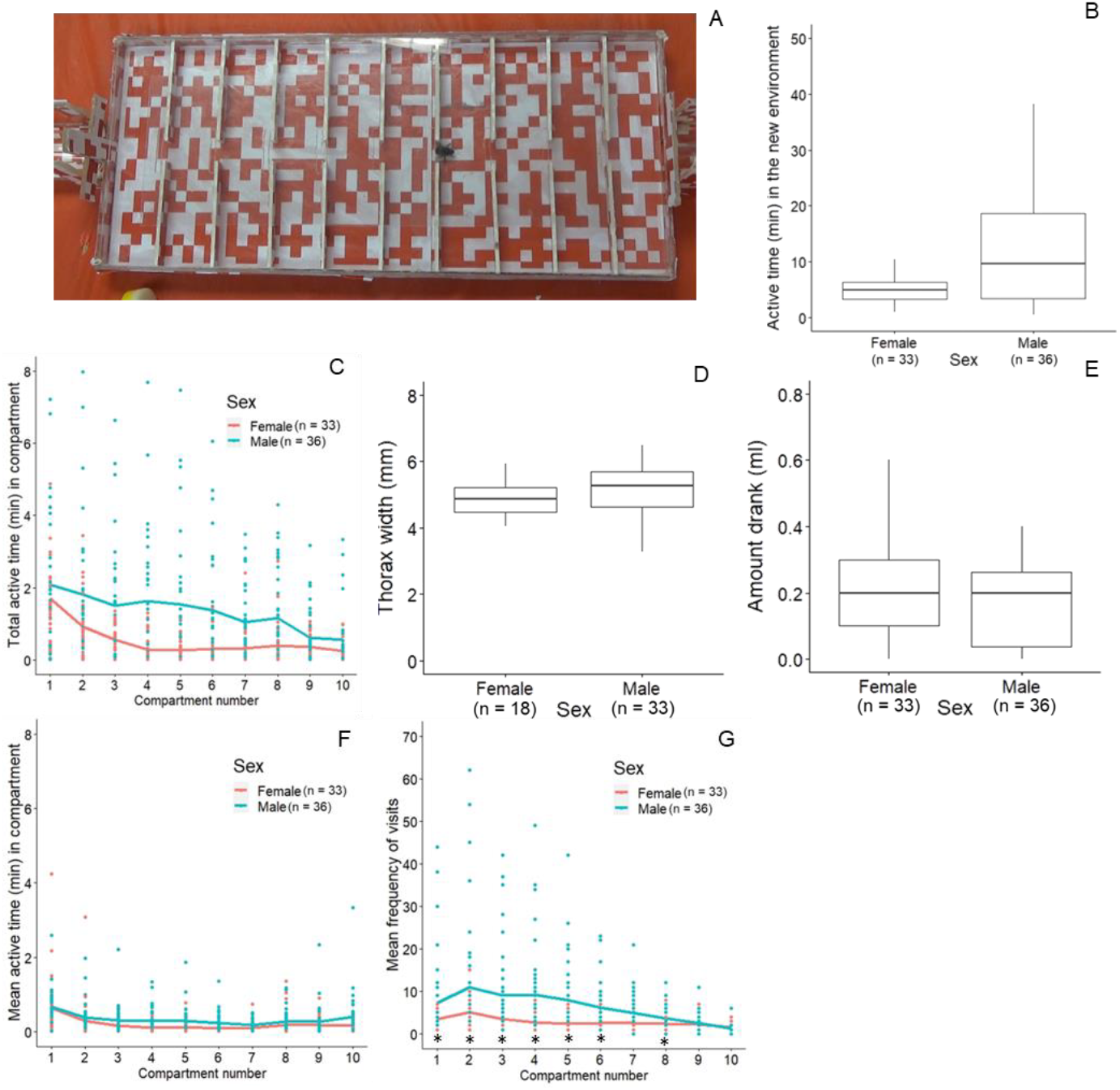
Results of the exploration task. **(A)** The exploration box had 10 equal-size compartments. Bees could freely move between the compartments through the holes in the middle of the dividers. **(B)** Distribution of the total active time (minutes) in all compartments for females and males. **(C)** Individual data (dots) and line graph showing the mean of total active time in minutes in each compartment for females (red) and males (blue). **(D)** Thorax width (mm) of females and males. **(E)** Sucrose consumption (ml) of females and males. **(F)** Individual data (dots) and line graph showing the mean active time in each compartment for females and males. **(G)** Individual data (dots) and mean (lines) frequencies of visiting each compartment females and males. * indicates *p*<0.05

As expected, the total active time in all compartments was significantly longer, and more variable in males (median = 11 minutes) than in females (median = 5 minutes) (GLMM: *p* = 0.012, figure 1B). Active time in each compartment was also significantly higher in males than females (*p* = 0.002, figure 1C). Active time significantly decreased across successive compartments (*p* < 0.001) but no interaction effect was detected between sex and compartment number (*p* = 0.899). Males and females who completed this task had a comparable body size (thorax width or inter-tegular span [35,36]) (*p* = 0.317, figure 1D) and sucrose consumption (*p* = 0.828, figure 1E). Within-sex analyses showed that, in males, the sucrose amount drank was not related to body size (*p* = 0.083) but was positively associated with active time (*p* = 0.008). In females, the sucrose amount drank was also not related to body size (*p* = 0.123) but was negatively associated with active time (*p* = 0.044). See full results in Table S1.

Males’ higher active time in each compartment may be driven by longer but fewer visits, equally frequent visits or shorter but more frequent visits in each compartment. We further analysed males’ strategy by examining sex differences in mean active time in each compartment (i.e., total active time in each compartment divided by the number of visits to that compartment) and the frequency of visiting each compartment. Bees significantly decreased the mean active time in each compartment (*p* < 0.001). Males and females had comparable mean active time in each compartment (*p* = 0.171). The interaction effect of sex and compartment number was not significant (*p* = 0.103, figure 1F). Males also visited each compartment more often than females (*p* < 0.001, figure 1G). The frequency of visits significantly decreased in successive compartments (*p* < 0.001) and there was a significant interaction between the frequency of visits and sex (*p* < 0.001). See full results in Table S1. Pairwise contrasts of sex differences in the frequency of visits to each compartment showed that males visited the compartments close to the entrance and in the middle of the box more often than females (Figure 1G, Table S2).

### Sex and Learning Performance of a Colour-Reward Association

The day after the exploration task, 29 bees (42%) re-emerged from the colony and participated in the discrimination-reversal colour learning task [29]. This task had a discrimination learning phase (DLP) followed by a reversal learning phase (RLP). Both learning phases were set in another box with 10 compartments of equal size (figure 2A, Note S3). Each compartment had two ‘flowers’, one blue and one yellow (each 4 cm x 4 cm), horizontally positioned on either side of the compartment, equidistant from the entrance (Note S4). Each bee was randomly assigned to associate one flower colour with a reward (0.01ml 50 w/w sucrose) and the other flower colour with a control (0.01ml water). Bees could explore both flowers, and their first choice (indicated by at least half of his/her body on a flower) was marked as either ‘correct’ (reward) or ‘incorrect/error’ (control). When a bee chose the rewarded flower, the shutter in the middle of the divider was lifted, and the bee passed to the next compartment. When the bee made an error, s/he had to visit the other flower (i.e., correctional choice) before going to the next compartment. A bee completed 1-2 sessions daily (as a measure of motivation and to avoid overfeeding). Each session had 10 flower pairs (i.e., 10 choices), and the side of the coloured flowers was pseudo-randomly determined for each trial. All bees experienced the same colour sequence in each trial to control for the order effect. The criterion for completing the DLP was ≥ 8 out of 10 rewarded choices as first choice (i.e., ≥80% correct) in two consecutive sessions.

**Figure 2.**
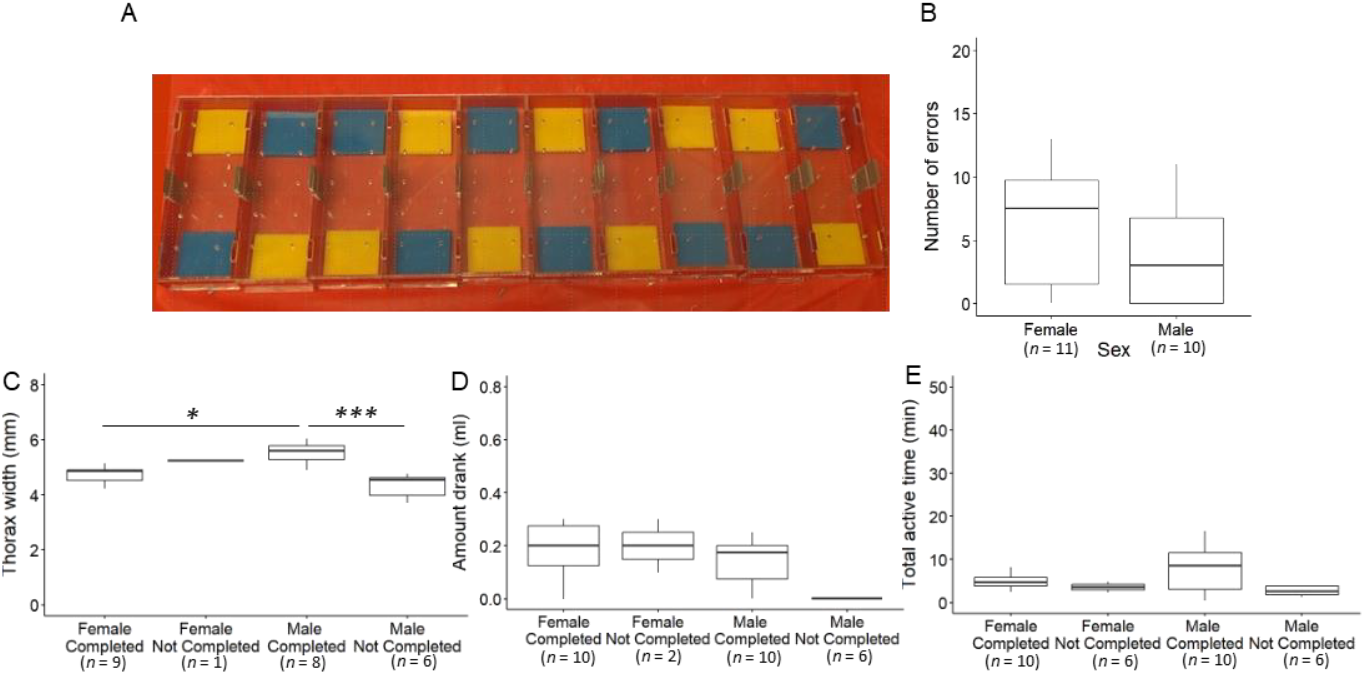
Results of the discrimination learning phase (DLP). **(A)** The discrimination-reversal learning task consisted of 10 equal-sized compartments. A shutter was in the middle of each divider, which was lifted after the bee made a correct choice (rewarded flower). **(B)** The number of errors (as a measure of associative learning performance) in the DLP for both sexes, including bees with the adjusted learning criterion. **(C-E)** Between-group pairwise comparisons between females and males that completed and did not complete the DLP. Note that sample size varies depending on the analyses. The ‘completed’ group included bees with the adjusted learning criterion whereas the ‘not-completed’ group included bees who participated in the learning phase but did not reach the learning criterion: **(C)** Thorax width (mm), **(D)** Sucrose consumption (ml), and **(E)** Total active time (mins) in the exploration task.* *p* < 0.05, *** *p* < 0.001.

Out of 29 bees, fifteen (8 females, 7 males) met the DLP learning criterion. The ability to learn the floral colour-reward association was measured as the number of errors made before reaching the DLP learning criterion [37–41]. Males made significantly fewer errors than females (*p* = 0.022) but this significance disappeared after including a few more bees that had stopped returning to the task but showed significant learning (adjusted criterion: a single session with 80% correct and ≥ 5 consecutive correct choices, *p* ≤ 0.031) (*n*_Female_ = 11, *n*_Male_ = 10, *p* = 0.139) (figure 2B).

Using the adjusted criterion, we further examined between-group differences (i.e., females and males who completed and did not complete the DLP) in body size, total active time and sucrose consumption during the exploration task using pairwise contrasts (tukey corrections for multiple comparisons). Full results are listed in table S3. Males of the completed group had a significantly larger body size than the females of the completed group (*p* = 0.042), and males of the not-complete group (*p* < 0.001) (figure 2C). The groups were not significantly different in sucrose consumption (figure 2D) and total active time in the exploration task (figure 2E). We also conducted within-sex analyses for the bees that had completed the task. Within-males analyses showed that none of the factors predicted males’ DLP performance (*n* = 10, body size: *p* = 0.298; amount drank: *p* = 0.079; active time: *p* = 0.482) whereas within-females analyses showed that the total active time was negatively related to errors in females (*n* = 9, body size: *p* = 0.304; amount drank: *p* = 0.163; active time: *p* = 0.021).

### Sex and Behavioural Flexibility

The 15 bees (8 females, 7 males) that had met the (unadjusted) learning criterion participated in the reversal learning phase (RLP) on the next day. The RLP had the same protocol and learning criterion as the DLP (detailed procedures in Note S4). During the RLP, the flower-reward contingency changed so that the bees had to unlearn the previous colour-reward association (e.g., B+ Y-), and relearn that the previously unrewarded colour became rewarded (e.g., B-Y+).

Thirteen bees (out of 15, all of the 8 females and 5 males) met the RLP learning criterion. Behavioural flexibility was measured from the number of errors made before reaching the learning criterion in the reversal learning phase (RLP) [37–41]. Overall, more errors were made before reaching the learning criterion in the RLP than in the DLP (*p* < 0.001) and males made significantly fewer errors than females (*p* = 0.013, figure 3A). No interaction effect was detected between sex and learning phase (*p* = 0.719). These results are retained after including the remaining two males who had stopped returning to the task but met the adjusted learning criterion of RLP (*p* = 0.023), or excluding a female who made extensive errors before reaching the learning criterion (*p* = 0.049).

**Figure 3.**
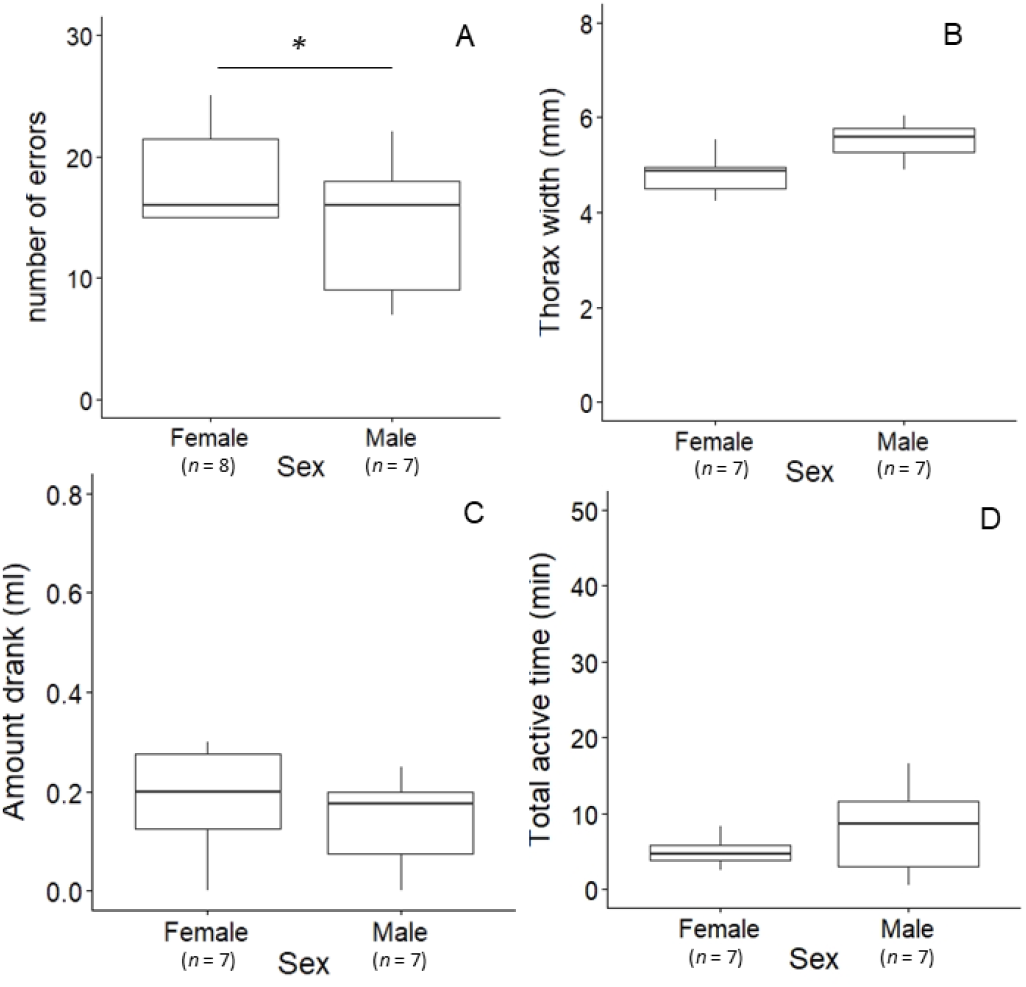
Results of the reversal learning phase (RLP). Between-group comparisons of females and males that completed the RLP including bees that met the adjusted criterion. (A) The number of errors (behavioural flexibility) in RLP for females and males. (B) thorax width (mm), (C) amount drank (ml) and (D) total active time (mins). The sample size decreased for the female group in B-D analysis due to data loss. * *p* < 0.05.

Using the adjusted learning criterion, all females and males completed the RLP (*n* = 15), and thus the between-group analyses included only females and males completed groups. No significant group differences in body size (*p* = 0.592, figure 3B), sucrose consumption (*p* = 0.210, figure 3C) and total active time during the exploration task (*p* = 0.266, figure 3D).

Within-sex analyses on the bees that had completed the RLP showed that none of these factors predicted males’ RLP performance (one male lost tag before thorax measure, *n* = 6, body size: *p* = 0.085; n = 7, amount drank: *p* = 0.093; active time: *p* = 0.335) whereas females (one bee lost her tag after completing the RLP, *n* = 7) who had a larger body size made more errors (body size: *p* < 0.001; amount drank: *p* = 0.430; active time: *p* = 0.596).

## Discussion

Exploration and behavioural flexibility may be expressed differently between sexes in animal species that have strong sex roles (e.g., [6–9]), such as bumblebees. We found evidence that compared with females, males are more exploratory (higher active time in a new environment) and show higher behavioural flexibility (made fewer errors before reaching the learning criterion in the reversal learning phase).

Males in general showed more exploration than females, and such a difference cannot be attributed to body size or sucrose consumption but is likely because drones generally fly longer than workers during dispersal [17–21]. During the exploration task, males overall visited most compartments more often than females. These results can be related to the field studies showing that males disperse in a random direction from their nest and travel far away from it [17,19–21], potentially increasing their likelihood of finding an unrelated queen for reproduction [22]. The frequency and mean duration of visits may also reflect the patrolling behaviour of males after dispersal; males create a flight path, marking their scent in different places to attract potential mates and regularly visit these places [42,43]. Within-sex analysis of males highlights the positive relationship between active time (exploration) and sucrose consumption, which may reflect that a higher active time leads to an increased energy expense, but may also suggest that males explore their environment more, which may increase their chances of discovering profitable flowers during dispersal [17].

Consistent with previous findings [23,25], males show comparable associative learning performance to females in the DLP. However, few bees have completed the DLP. Among the females that completed the DLP, the active time (exploration) was correlated to faster associative learning performances, suggesting that the role of exploration is important for workers [45]; a strong association between exploration tendency and associative learning has been shown to benefit the colony when competing to efficiently exploit newly emerged food sources [44]. It is assumed that exploration plays a potential role in males’ foraging.

However, within-males analysis showed no relationship between associative learning and exploration as well as other tested factors (i.e., body size, sucrose consumption). Despite this, the between-group analysis showed that males who completed the DLP had a larger body size than females who had completed the DLP and the males who never reached the DLP learning criterion, suggesting that a larger body size may bring an advantage for males. Overall, the results of DLP are in line with previous studies that used different setups (e.g., stimuli presented in an array or two-choice task) and conditions (e.g., natural or laboratory) [23,25]. However, further work with these setups should be conducted to clarify the potential link between bees’ tendencies for exploration and their associative learning abilities and increase the generalisability of our own results.

In the RLP, bumblebees made more errors than in the DLP, suggesting that the previously rewarded stimulus may become a ‘distractor’ when they had to relearn the reward contingency [27]. As predicted, males were more behaviourally flexible (i.e., made fewer errors) than females. Because females and males that completed the RLP had comparable body size, sucrose consumption and total active time in the exploration task, these factors cannot explain the sex difference in behavioural flexibility. Unlike females where body size is a predictor for their RLP performance, within-males analysis showed that none of the tested factors predict males’ RLP performance, suggesting that males’ behavioural flexibility can be independent of other factors. The correlation between workers’ size and performance in the RLP may be due to a difference in foraging roles between large and small workers.

Indeed, previous work has shown that larger bees invest more in learning highly rewarding flowers (like in our experiment) while small bees more readily accept flowers of all qualities [26]. On the other hand, the overall enhanced flexibility of males can be explained by their readiness to adopt the ‘win-stay, lose-shift’ strategy in the face of a change. This strategy is ecologically adaptive (e.g., in response to the change in flower quality [28]), which can increase foraging efficiency and maximise energy gain while searching for mates for reproduction. Compared with males, the costs of not being flexible are lower for females who can continue to access resources in the colony. There is also an upside for workers making more ‘mistakes’, which can lead to discovering and exploiting new profitable sources and collecting more resources for the colony [46].

In conclusion, we provided evidence that exploration and behavioural flexibility are two key behavioural and cognitive traits for male bumblebees in the context of their dispersal. Drones outperform workers in exploration and behavioural flexibility. The enhanced exploration and behavioural flexibility could be related to the pressure of being solitary foraging when searching for mates for reproduction. The enhancement in these two traits likely brings advantages to males in mating and foraging contexts. These results overall contribute to the understanding of how the expression of traits that have ecological and evolutionary importance on fitness vary between sexes with different roles.

## Author contributions

PKYC designed, conducted, analysed, and wrote the first draft of the study. SD conducted the experiment and preliminary analyses. KH and TR made significant contributions to conceiving the project idea, analyzing it, and discussing it.

## Acknowledgment

We thank Toyota Manufacturing Deeside for providing full support for conducting this experiment. Thanks also to M MacDonald, O Reece, L Chumper, and beekeepers E Donnelly and B Donnelly for their logistical help.

## Pre-print

This study has been submitted to BioRxiv.

## Supplementary materials

### Note S1. *Study species and housing*

We conducted this experiment between May and August 2024. Bumblebees from 3 colonies and 2 pocket hives purchased from commercial suppliers were used in the study (BIOBEST from Biobest Belgium N.V., Westerlo, Belgium, and Koppert, UK). Two colonies were housed in a natural environment based in Toyota Manufacturing Deeside and later brought to the lab for the study when there were few individuals left in the colony. The other colony and pocket hives were housed in the lab. The lab setup mirrored the bees’ natural housing as much as possible. The lab (room temperature 21-23^°^c) was lit with natural daylight, with additional artificial lights provided for the bees. Bees were kept in a black box mirroring their underground nest. We extended the housing structure by attaching a transparent circular (diameter 4 cm) tunnel to a pocket hive or a colony. The tunnel had three shutters and an open end where a plunger could be placed to capture the bee during the experiment. Bees were individually marked with a number using shellac-made glue [30].

### Note S2. *Ethics*

Animal welfare and husbandry were carried out daily whereby bees were fed syrup and pollen. The bees had *ad libitum* access to syrup and pollen in the exploration task but we decreased the amount of syrup and pollen to them during the learning task to increase their motivation. Enrichments (e.g., paper sticks and tissue tiles) were provided for the bees. While there is no general animal welfare and husbandry guideline for invertebrates, this study strictly followed ASAB and ABS animal welfare guidelines and was approved by the Division of Psychology Ethics Committees (no: SDPC130624).

### Note S3. *Apparatus*

The experiment used two apparatuses (figure 1 and figure 3). Both boxes were designed for bees to mostly walk to the stimuli. One apparatus was made for the exploration task (Height x Width x Length: 3.3 cm x 21.8 cm x 50.9 cm, figure 1A) and the other one was made for the discrimination-reversal learning task (H x W x L: 3.1 × 15.6 × 53.6 cm, figure 3A). Both apparatuses shared similar features; both boxes were rectangular-shaped and had 10 equally divided cells. Each cell had a hole in the middle of the wall separating it from the next one and a shutter was used to block this entrance between compartments, and at the one end of the apparatus. This allowed the experimenter (PC and SD) to lift the shutter and let a bee enter/exit the experiment or go to the next compartment through the holes. In the exploration apparatus, the floor of each cell (W x H: each 4.5 × 1.2 cm) was covered with a random mosaic white-and-red square (1 cm x 1 cm) pattern. In the learning apparatus, each cell had a pair of colour stimuli (each 4 cm x 4 cm). The colour and size of the stimuli were chosen based on previous studies,^e.g.,[47]^ yellow (Perspex® Yellow 250) and blue (Perspex® Blue 727). The stimuli were horizontally positioned, one on the left side and one on the right side of the compartment, equidistant from the entrance.

### Note S4. Procedure

All the bees went through the same standardised procedure, they left home the first time to participate in the exploration task. When a bee emerged from its colony and went to the end of the tunnel, s/he was brought to the task with a plunger placed at the end of the tunnel.

The exploration task was designed to measure exploration and activity levels in a novel environment. The task started when a bee’s full body entered the exploration box and ended when the bee’s full body exited the box. If a bee was inactive for 15 minutes in the box, the session was terminated and the bee was brought home. If the bee completed the session, s/he was fed with 50 w/w sucrose *ad libitum* (an optimal food source preferred by both sexes) [32–34]; this aimed to assess whether they were responsive to sucrose, and the drinking amount was measured.

After the exploration task, the bees had an additional two training sessions that served two purposes; 1) to allow the bees to familiarise themselves with the procedure of the learning task, and 2) to ensure the bees had high motivation for the learning task. In these training sessions, a shutter was added to block the hole so that the bees had to explore both the right and left sides of the box. The shutter between the compartment and the next one was lifted every 30 seconds and the bees had to go through all the 10 compartments to be considered as passing the training phase. After each training, the bees were rewarded with 50 w/w sucrose.

The bees that had participated in the exploration task and passed the training phases went to the learning task (full completion as an indicator of motivation). The learning task consisted of a discrimination learning phase (DLP) followed by a reversal learning phase (RLP).

During each ‘session’, the bees had to choose between two colours in each compartment (a total of 10 choices). In the DLP, one of the flower colours, randomly assigned to each bee, was associated with a favourable reward (e.g., B+ *or* Y+), a drop of 0.01ml 50 w/w sucrose at the centre of the flower. The other flower colour had a drop of 0.01ml water (control), also placed at the centre of the flower. The bees could explore both flowers, and their first choice (indicated as when at least half of his/her body was on a flower) was marked as either ‘correct’ (the rewarded flower) or ‘incorrect’ (the control). When they made an ‘incorrect’ choice (or error), they had to visit the other flower (i.e., correctional choice) before passing to the next compartment through the hole in the middle of the side wall. Choices were recorded both by direct observation and a video camera (Panasonic) facing downward 1.5 m above the apparatus and attached to a tripod 30 cm away from the box.

A bee went through 1-2 sessions daily (as a motivation measure), with each session including 10 flower pairs (i.e., made 10 choices per session). The presentation of the flower pairs was pseudo-randomised within and across sessions. Within sessions, the bees saw the same flower colour on the left side five times and on the right side five times. The same flower colour was presented on the same side for no more than two consecutive compartments to avoid bees developing side bias. Across sessions, the same colour pair was not presented in the same compartment for more than two consecutive times. All the bees experienced the same colour sequence to control the order effect on the experience. The bee was brought back home when s/he completed the task and went to the second session if they emerged from the colony 40 minutes or more after the last session. After each session, the box was cleaned with water to remove any scent left from the previous bee and reset for the next bee. The criterion for completing this learning phase was when a bee chose the rewarded flower as the first choice at least 8 out of 10 times (80%) in two consecutive sessions. Learning ability was measured as the number of errors made before reaching the criterion.

Bees went to the reversal learning phase (RLP) the day after they had completed the DLP. The RLP used the same protocol and learning criterion as in the DLP. However, during RLP, the bees had to learn that the previously rewarded colour was no longer rewarded (e.g., B+ Y-), and that the previously unrewarded colour was now rewarded (e.g., B-Y+).

### Note S5. Behavioural measurement

Active time in the compartment (a proxy of exploration) was measured during which the bee experienced the novel apparatus for the first time. The recording started when a bee’s full body entered the apparatus (the first compartment) until its full body left the apparatus (the last compartment). From these recordings, we obtained the active time of each bee in each compartment and the total time in all 10 compartments.

The performance of associative learning and behavioural flexibility was the number of errors made before reaching the learning criterion in each learning phase (DLP and RLP). The stringent learning criterion was 80% correct responses for two consecutive sessions.

Associative learning performance was the number of errors made before reaching the learning criterion in the DLP and behavioural flexibility performance was the number of errors made before reaching the learning criterion in the RLP. We also adjusted the learning criterion to include bees that had not returned to the task but showed significant learning, which was a bee met 80% correct responses in a session and that there were 5 consecutive correct responses or more in that session.

The size of each bee was estimated using the thorax width or inter-tegular span [35,36]. The sucrose consumption (ml) was the amount consumed after exploration using a syringe with a 0.02 ml interval.

### Note S6. Statistical analyses

All analyses were conducted using R (version 4.4.1) and the significance level was set as two-tailed *p* ≤ 0.05. For the exploration task, a Generalised Linear Mixed Model (GLMM) with gamma log link distribution in the ‘glmmTMB’ package [48] was used to examine 1) whether sex and body size predicted the total active time in all the compartments; 2) sex, compartment number, and their interaction in relation to the total active time in each compartment; 3) sex, compartment number, and their interaction in relation to the mean active time in each compartment; and 4) sex difference in body size. In all GLMM models, we included bee ID as a random variable to maximise model convergence. Beta regression of the ‘betareg’ package [49] was used to examine whether sex, body size, and the total active time in all the compartments predicted sucrose consumption. Poisson log link distribution was used to determine the frequency of visiting each compartment in the exploration task, the number of errors made in the DLP, and the number of errors made in the RLP. Multicollinearity was checked after running each model using Variance Inflation Factor (VIF) (<5) and tolerance (0.25).

For each learning phase, two models were run to examine learning performance. In the first model, we only included bees that had met the learning criterion (80% correct responses for two consecutive sessions). This model included a fixed factor, sex, and the response variable was the number of errors made before reaching the learning criterion. We then included bees that had met the adjusted learning criterion (i.e., met the 80% learning criterion with 5 or more consecutive correct responses in a session) and reran the model as the second model. Both models included bee identity as random variables for model convergence purposes.

We then carried out within-sex analyses for bees that had completed each learning phase. We used GLMM Poisson log link distribution to examine whether their body size, sucrose consumption, and total active time in all compartments predicted their associative learning performance in one model and behavioural flexibility in another model. Both models included bee identity as the random variable.

Between-group analyses were conducted to examine whether females and males who had and had not completed each learning phase differed in body size, sucrose consumption, and total active time in the new environment. We first ran a GLMM for each analysis followed by pairwise contrasts with Tukey corrections for multiple comparisons using the ‘emmeans’ package [50]. GLMMs with gamma log link distribution were used to examine body size in one model and total active time in all compartments in another model. All GLMMs included bee identity as the random variable. Finally, a beta regression was used to examine sex differences in sucrose consumption.

**Table S1.** This table shows fixed variables, estimates (est), standard error (S.E), *z* and *p* values for response variable (a) total active time, (b) active time in each compartment (c) mean active time in each compartment and (d) frequency to each compartment. It also included within-sex analysis of (g) males and (h) females on their sucrose drinking amount. Bold values indicate significance *p* <0.05.

| | Response variable | Fixed variable(s) | Est | S.E | $z$ | $p$ |
| --- | --- | --- | --- | --- | --- | --- |
| a | Total active time | Sex (Male) | 1.06 | 0.41 | 2.56 | **0.011** |
|  |  | Thorax | 0.20 | 0.26 | 0.76 | 0.445 |
| b | Active time in each compartment | Sex (Male) | 0.90 | 0.30 | 3.03 | **0.002** |
|  |  | Compartment number | -0.20 | 0.02 | -10.14 | **< 0.001** |
|  |  | Sex (Male)* compartment number | < 0.01 | 0.03 | 0.13 | 0.899 |
| c | Mean active time in each compartment | Sex (Male) | 0.26 | 0.19 | 1.37 | 0.171 |
|  |  | Compartment number | -0.12 | 0.02 | -7.05 | **< 0.001** |
|  |  | Sex (Male)* compartment number | 0.04 | 0.02 | 1.63 | 0.103 |
| d | Frequency to each compartment | Sex (Male) | 0.80 | 0.20 | 4.00 | **< 0.001** |
|  |  | Compartment number | -0.10 | 0.01 | -8.17 | **< 0.001** |
|  |  | Sex (Male)* compartment number | -0.05 | 0.01 | 0.01 | **< 0.001** |
| e | Thorax size | Sex (Male) | 0.04 | 0.04 | 1.00 | 0.317 |
| f | Amount drank | Sex (Male) | -0.06 | 0.25 | -0.22 | 0.828 |
| Within-sex analysis |  |  |  |  |  |  |
| g | Males: | Thorax size | 0.41 | 0.23 | 1.73 | 0.083 |
|  | Amount drank | Active time | 0.01 | 0.01 | 2.65 | **0.008** |
| h | Females: | Thorax size | 0.66 | 0.43 | 1.54 | 0.123 |
|  | Amount drank | Active time | -0.13 | 0.07 | -2.02 | **0.044** |

**Table S2.** Pairwise contrasts of males’ and females’ frequent of visiting each compartment. This table shows estimates, standard error (S.E), *z* and *p* values. Bold values indicate significance *p* <0.05

| | Estimate | S. E | $z$ | $p$ |
| --- | --- | --- | --- | --- |
| Comp 1: Female - Male | -0.45 | 0.22 | -2.09 | **0.037** |
| Comp 2: Female - Male | -0.52 | 0.21 | -2.49 | **0.013** |
| Comp. 3: Female - Male | -0.69 | 0.22 | -3.20 | **0.001** |
| Comp. 4: Female - Male | -0.98 | 0.22 | -4.41 | **<0.001** |
| Comp. 5: Female - Male | -0.95 | 0.23 | -4.20 | **<0.001** |
| Comp. 6: Female - Male | -0.64 | 0.23 | -2.83 | **0.005** |
| Comp. 7: Female - Male | -0.40 | 0.23 | -1.77 | 0.077 |
| Comp. 8: Female - Male | -0.17 | 0.24 | -0.70 | **0.481** |
| Comp. 9: Female - Male | 0.13 | 0.25 | 0.52 | 0.604 |
| Comp. 10: Female - Male | 0.39 | 0.28 | 1.43 | 0.152 |

**Table S3.**
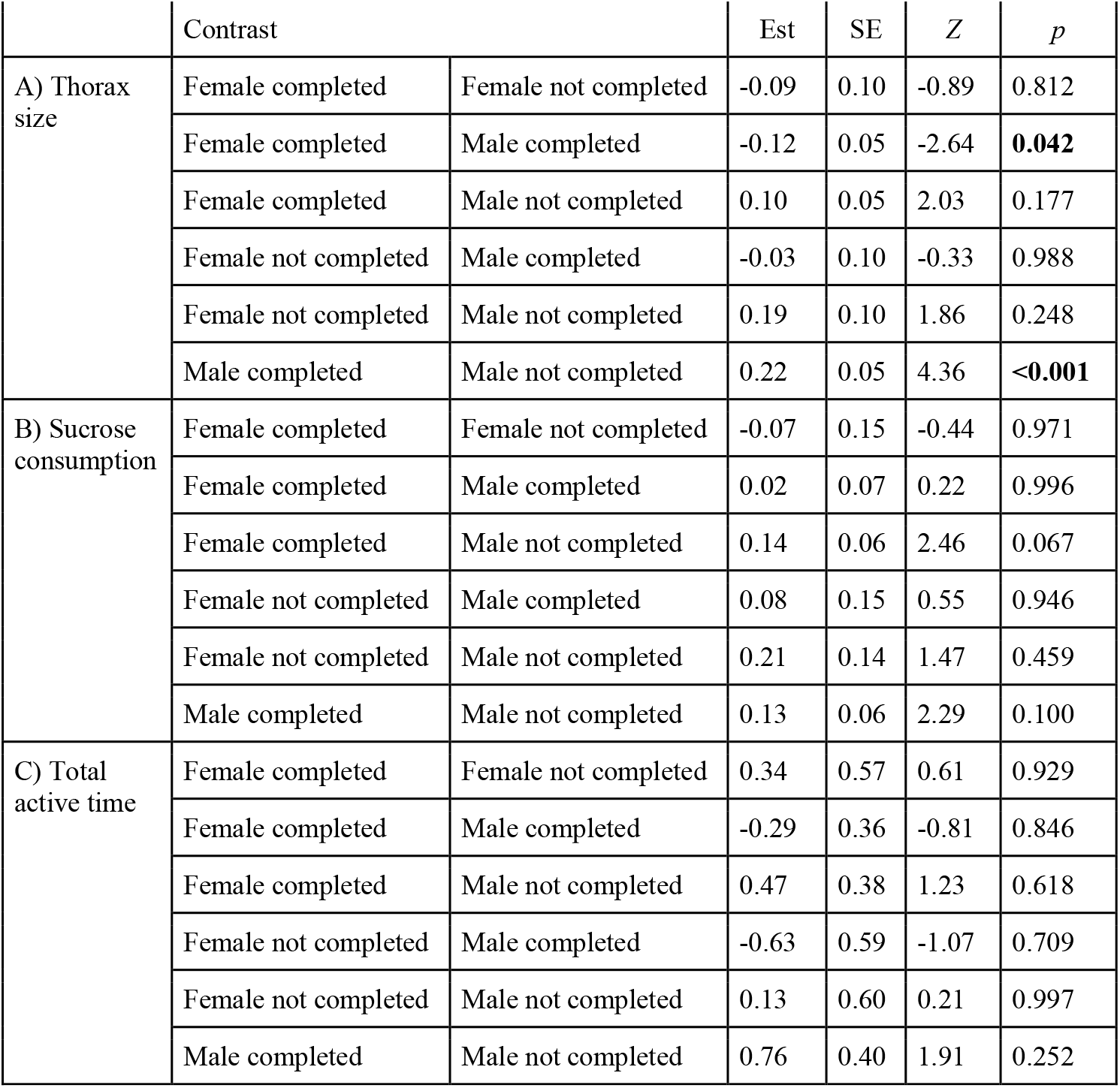
Pairwise contrast with tukey adjusted *p* values between females and males that completed the DLP and males that did not complete the DLP in A) thorax size, B) sucrose consumption, and C) total active time in the exploration task. This table shows the contrast groups, estimates, standard errors (S.E.), *Z* and *p* values. Bold values indicate *p* < 0.05.

| | Contrast | | Est | SE | $Z$ | $p$ |
| --- | --- | --- | --- | --- | --- | --- |
| A) Thorax size | Female completed | Female not completed | -0.09 | 0.10 | -0.89 | 0.812 |
|  | Female completed | Male completed | -0.12 | 0.05 | -2.64 | **0.042** |
|  | Female completed | Male not completed | 0.10 | 0.05 | 2.03 | 0.177 |
|  | Female not completed | Male completed | -0.03 | 0.10 | -0.33 | 0.988 |
|  | Female not completed | Male not completed | 0.19 | 0.10 | 1.86 | 0.248 |
|  | Male completed | Male not completed | 0.22 | 0.05 | 4.36 | **<0.001** |
| B) Sucrose consumption | Female completed | Female not completed | -0.07 | 0.15 | -0.44 | 0.971 |
|  | Female completed | Male completed | 0.02 | 0.07 | 0.22 | 0.996 |
|  | Female completed | Male not completed | 0.14 | 0.06 | 2.46 | 0.067 |
|  | Female not completed | Male completed | 0.08 | 0.15 | 0.55 | 0.946 |
|  | Female not completed | Male not completed | 0.21 | 0.14 | 1.47 | 0.459 |
|  | Male completed | Male not completed | 0.13 | 0.06 | 2.29 | 0.100 |
| C) Total active time | Female completed | Female not completed | 0.34 | 0.57 | 0.61 | 0.929 |
|  | Female completed | Male completed | -0.29 | 0.36 | -0.81 | 0.846 |
|  | Female completed | Male not completed | 0.47 | 0.38 | 1.23 | 0.618 |
|  | Female not completed | Male completed | -0.63 | 0.59 | -1.07 | 0.709 |
|  | Female not completed | Male not completed | 0.13 | 0.60 | 0.21 | 0.997 |
|  | Male completed | Male not completed | 0.76 | 0.40 | 1.91 | 0.252 |

## References

Réale D, Reader SM, Sol D, McDougall PT, Dingemanse NJ. 2007 Integrating animal temperament within ecology and evolution. Biol. Rev. Camb. Philos. Soc. 82, 291–318.

Dukas R. 2013 Effects of learning on evolution: robustness, innovation and speciation. Anim. Behav. 85, 1023–1030.

Reader SM. 2015 Causes of Individual Differences in Animal Exploration and Search. Top. Cogn. Sci. 7, 451–468.

Dukas R, Bernays EA. 2000 Learning improves growth rate in grasshoppers. Proc. Natl. Acad. Sci. U. S. A. 97, 2637–2640.

Nicolakakis N, Sol D, Lefebvre L. 2003 Behavioural flexibility predicts species richness in birds, but not extinction risk. Anim. Behav. 65, 445–452.

Videlier M, Cornette R, Bonneaud C, Herrel A. 2015 Sexual differences in exploration behavior in Xenopus tropicalis? J. Exp. Biol. 218, 1733–1739.

Ogurtsov SV, Antipov VA, Permyakov MG. 2018 Sex differences in exploratory behaviour of the common toad, Bufo bufo. Ethol. Ecol. Evol. 30, 543–568.

Lucon-Xiccato T, Bisazza A. 2014 Discrimination reversal learning reveals greater female behavioural flexibility in guppies. Biol. Lett. 10, 20140206.

Fuss T, Witte K. 2019 Sex differences in color discrimination and serial reversal learning in mollies and guppies. Curr. Zool. 65, 323–332.

Thomson JD, Plowright RC. 1980 Pollen carryover, nectar rewards, and pollinator behavior with special reference to Diervilla lonicera. Oecologia 46, 68–74.

Yadav S, Yadav S, Sharma D, Sangwan N. 2016 Bumblebees, Life Cycle and their role in Pollination-A Review. Int J Res Sci

Baer B. 2003 Bumblebees as model organisms to study male sexual selection in social insects. Behav. Ecol. Sociobiol. 54, 521–533.

Belsky JE, Camp AA, Lehmann DM. 2020 The Importance of Males to Bumble Bee (Bombus Species) Nest Development and Colony Viability. Insects 11. (doi:10.3390/insects11080506)

Goulson D. 2003 Bumblebees: Their Behaviour and Ecology. Oxford University Press.

Zhao H, Mashilingi SK, Liu Y, An J. 2021 Factors influencing the reproductive ability of male bees: Current knowledge and further directions. Insects 12, 529.

Robert T, Frasnelli E, Collett TS, Hempel de Ibarra N. 2017 Male bumblebees perform learning flights on leaving a flower but not when leaving their nest. J. Exp. Biol. 220, 930–937.

Kraus FB, Wolf S, Moritz RFA. 2009 Male flight distance and population substructure in the bumblebee Bombus terrestris. J. Anim. Ecol. 78, 247–252.

Bertsch A. 1984 Foraging in male bumblebees (Bombus lucorum L.): maximizing energy or minimizing water load? Oecologia 62, 325–336.

Wolf S, Toev T, Moritz RLV, Moritz RFA. 2012 Spatial and temporal dynamics of the male effective population size in bumblebees (Hymenoptera: Apidae). Popul. Ecol. 54, 115–124.

Osborne JL, Clark SJ, Morris RJ, Williams IH, Riley JR, Smith AD, Reynolds DR, Edwards AS. 1999 A landscape-scale study of bumble bee foraging range and constancy, using harmonic radar. J. Appl. Ecol. 36, 519–533.

Chapman RE, Wang J, Bourke AFG. 2003 Genetic analysis of spatial foraging patterns and resource sharing in bumble bee pollinators. Mol. Ecol. 12, 2801–2808.

Amin MR, Bussière LF, Goulson D. 2012 Effects of Male age and Size on Mating Success in the Bumblebee Bombus terrestris. J. Insect Behav. 25, 362–374.

Wolf S, Chittka L. 2016 Male bumblebees, Bombus terrestris, perform equally well as workers in a serial colour-learning task. Anim. Behav. 111, 147–155.

Manning TH, Austin MW, MuseMorris K, Dunlap AS. 2021 Equivalent learning, but unequal participation: Male bumble bees learn comparably to females, but participate in cognitive assessments at lower rates. Behav. Processes 193, 104528.

Muth F, Tripodi AD, Bonilla R, Strange JP, Leonard AS. 2021 No sex differences in learning in wild bumblebees. Behav. Ecol. 32, 638–645.

Frasnelli E, Robert T, Chow PKY, Scales B, Gibson S, Manning N, Philippides AO, Collett TS, Hempel de Ibarra N. 2021 Small and Large Bumblebees Invest Differently when Learning about Flowers. Curr. Biol. 31, 1058–1064.e3.

Dyer AG, Chittka L. 2004 Fine colour discrimination requires differential conditioning in bumblebees. Naturwissenschaften 91, 224–227.

Wolf S, Moritz RFA. 2014 The pollination potential of free-foraging bumblebee (Bombus spp.) males (Hymenoptera: Apidae). Apidologie 45, 440–450.

Shettleworth SJ. 2010 Cognition, Evolution, and Behavior. Oxford University Press.

Toppa RH, Arena MVN, da Silva CI, Marendy P, de Souza P, da Silva-Zacarin ECM. 2021 Impact of glues used for RFIDs on the longevity and flight muscles of the stingless bee Melipona quadrifasciata (Apidae: Meliponini). Apidologie 52, 328–340.

Wu NC, Seebacher F. 2022 Physiology can predict animal activity, exploration, and dispersal. Commun. Biol. 5, 109.

Brown M, Brown MJF. 2020 Nectar preferences in male bumblebees. Insectes Soc. 67, 221–228.

Bailes EJ, Pattrick JG, Glover BJ. 2018 An analysis of the energetic reward offered by field bean (Vicia faba) flowers: Nectar, pollen, and operative force. Ecol. Evol. 8, 3161– 3171.

Pamminger T, Becker R, Himmelreich S, Schneider CW, Bergtold M. 2019 The nectar report: quantitative review of nectar sugar concentrations offered by bee visited flowers in agricultural and non-agricultural landscapes. PeerJ 7, e6329.

Hagen M, Dupont YL. 2013 Inter-tegular span and head width as estimators of fresh and dry body mass in bumblebees (Bombus spp.). Insectes Soc. 60, 251–257.

Cane JH. 1987 Estimation of Bee Size Using Intertegular Span (Apoidea). J. Kans. Entomol. Soc. 60, 145–147.

Izquierdo A, Jentsch JD. 2012 Reversal learning as a measure of impulsive and compulsive behavior in addictions. Psychopharmacology 219, 607–620.

Wascher CAF, Allen K, Szipl G. 2021 Learning and motor inhibitory control in crows and domestic chickens. R Soc Open Sci 8, 210504.

Tapp PD, Siwak CT, Estrada J, Head E, Muggenburg BA, Cotman CW, Milgram NW. 2003 Size and reversal learning in the beagle dog as a measure of executive function and inhibitory control in aging. Learn. Mem. 10, 64–73.

Nilsson SRO, Alsiö J, Somerville EM, Clifton PG. 2015 The rat’s not for turning: Dissociating the psychological components of cognitive inflexibility. Neurosci. Biobehav. Rev. 56, 1–14.

Giurfa M, Hammer M, Stach S, Stollhoff N, Müller-deisig N, Mizyrycki C. 1999 Pattern learning by honeybees: conditioning procedure and recognition strategy. Anim. Behav. 57, 315–324.

Valterová I, Martinet B, Michez D, Rasmont P, Brasero N. 2019 Sexual attraction: a review of bumblebee male pheromones. Z. Naturforsch. C 74, 233–250.

Alcock J, Barrows EM, Gordh G, Hubbard LJ, Kirkendall L, Pyle DW, Ponder TL, Zalom FG. 1978 The ecology and evolution of male reproductive behaviour in the bees and wasps. Zool. J. Linn. Soc. 64, 293–326.

Raine NE, Chittka L. 2008 The correlation of learning speed and natural foraging success in bumble-bees. Proc. Biol. Sci. 275, 803–808.

Dougherty LR, Guillette LM. 2018 Linking personality and cognition: a meta-analysis. Philos. Trans. R. Soc. Lond. B Biol. Sci. 373, 20170282.

Evans LJ, Smith KE, Raine NE. 2017 Fast learning in free-foraging bumble bees is negatively correlated with lifetime resource collection. Sci. Rep. 7, 496.

Strang CG, Sherry DF. 2014 Serial reversal learning in bumblebees (Bombus impatiens). Anim. Cogn. 17, 723–734.

Magnusson A, Skaug H, Nielsen A, Berg C, Kristensen K, Maechler M, van Bentham K, Bolker B, Brooks ME. 2017 glmmTMB: generalized linear mixed models using template model builder. R package version 0. 1 3.

Cribari-Neto F, Zeileis A. 2010 Beta Regression in R. Journal of Statistical Software, Articles 34, 1–24.

Lenth, Singmann, Love, Buerkner, Herve. In press. Emmeans: Estimated marginal means, aka least-squares means. R package version

